# *Pseudomonas aeruginosa* partitioning protein ParB acts as a nucleoid-associated protein binding to multiple copies of a *parS*-related motif

**DOI:** 10.1101/280743

**Authors:** Adam Kawalek, Aneta Agnieszka Bartosik, Krzysztof Glabski, Grazyna Jagura-Burdzy

**Affiliations:** Institute of Biochemistry and Biophysics, Polish Academy of Sciences, Department of Microbial Biochemistry, Pawinskiego 5a, 02-106 Warsaw, Poland

## Abstract

ParA and ParB homologs are involved in accurate chromosome segregation in bacteria. ParBs participate in separation of ori domains by binding to specific *parS* sites, mainly localized close to *oriC*. In *Pseudomonas aeruginosa* neither a lack of *parB* gene nor modification of ten *parS*s is lethal. Remarkably, such mutants show not only defects in chromosome segregation but also growth retardation and motility dysfunctions. Moreover, a lack of *parB* alters expression of over one thousand genes, suggesting that ParB could interact with the chromosome outside its canonical *parS* targets.

Indeed, DNA immunoprecipitation with anti-ParB antibodies followed by deep sequencing (ChIP-seq) revealed 420 enriched regions in WT PAO1161 strain and around 1000 in a ParB-overproducing strain and in various *parS* mutants. Vast majority of the ParB-enriched loci contained a heptanucleotide motif corresponding to one arm of the *parS* palindrome. All previously postulated *parS* sites with the exception of *parS5* interacted with ParB *in vivo.* Whereas the ParB binding to the four *parS* sites closest to *oriC, parS1-4*, is involved in chromosome segregation, its genome-wide interactions with hundreds of *parS* half-sites could affect chromosome topology, compaction and gene expression classifying *P. aeruginosa* ParB as a Nucleoid Associated Protein (NAP).

## INTRODUCTION

Bacterial genome segregation is a precise, complex and still only partially understood process. Our knowledge of it originates from studies on low-copy-number plasmids in which partitioning systems have been identified (1, 2). Despite the diversity of these systems, they all contain two protein components: A, a NTPase securing energy supply and B, a DNA binding protein, and the third vital element, a specific DNA motif, designated centromere-like sequence (*parC/parS*), recognized and bound by B component (2–4). The discovery of a *parA-parB* operon located close to *oriC* and encoding homologs of plasmid partitioning proteins of class IA in the majority of the bacterial chromosomes sequenced, has implicated its potential role in chromosome segregation (5–8).

Most of the information obtained by studying the segregation of low-copy-number plasmids seems to be relevant to the structure and action of the chromosomal homologs, despite their species-specific scope of importance. Deletion of *parA-parB* genes is lethal in some species, e.g., *Caulobacter crescentus* or *Myxococcus Xanthus* (9, 10) whereas in other species *par* mutants demonstrate defects of chromosome segregation of various severity in the vegetative and/or sporulation phase of growth, e.g., *Bacillus subtilis*, *Streptomyces coelicolor*, *Mycobacterium smegmatis*, *Vibrio cholerae*, *Streptococcus pneumoniae*, *Corynebacterium glutamicum*, *Pseudomonas aeruginosa* (11–25). Chromosomal ParAs are Walker-type ATPases lacking the specific DNA binding domain present in the N-terminal part of the plasmid ParAs (26, 27). Chromosomal ParBs, similarly to the plasmidic counterparts, contain an extended HTH motif, a C-terminal dimerization domain and an N-terminal polymerization domain (4, 28–34) and have an ability to spread around the *parS* sequences (34–37) and silence nearby genes, at least in test plasmids (31).

Chromosomal ParBs recognize multiple *parS* sequences (2–20), highly conserved across the bacterial kingdom and localized mainly in the ori domains of primary chromosomes (defined as 20% around the origin of replication *oriC*) (8, 28, 38). They form large nucleoprotein complexes organizing newly replicated ori domains and, with the help of ParAs, separate these domains and hold them in defined locations near the poles until the end of replication and cytokinesis (12, 18, 39– 45). It has been shown that in *B. subtilis* and *S. pneumoniae* ParB-bound *parS* sites act as loading platforms for SMC complexes facilitating condensation of ori domains and subsequently all newly replicated DNA (21, 46–50). Two major hypotheses of how the ParBs of Class IA build the large nucleoprotein complexes around the *parS* sequences have been put forward. Both are based on the data that ParBs bind to palindromic *parS* sequences as dimers but can also laterally spread along DNA due to dimer-dimer interactions and weak interactions with nonspecific DNA. In the looping and bridging model ParB dimers bound to DNA interact at a distance, which leads to further DNA compaction (37, 51). In the nucleation and caging model ParB bound to *parS* acts as a nucleation center in which protein-protein and protein-nsDNA interactions spatially entrap additional ParB molecules forming a large nucleoprotein complex (52).

Besides their involvement in genome segregation chromosomal ParAs and ParBs have also been shown to influence bacterial growth and cell cycle by affecting DNA replication and cell division, competence or motility (14, 15, 20, 23, 24, 39, 40, 53).

In turn, in addition to their main function in plasmid partitioning, plasmidic ParBs of class IA may act as global transcriptional regulators by binding to their operators, spreading and silencing nearby promoters e.g., in the IncP-1 plasmids (54, 55) or IncU plasmids (56). In contrast, under native conditions oligomers of chromosomal ParBs spread by up to 20 kb from the *parS*s (35–37, 51) but seem to affect gene expression very selectively. A comparison of the transcriptomes of *B. subtilis* WT strain and *spo0J*_null_ mutant (Spo0J, a homolog of ParB) did not reveal significant changes around ten *parS* sites (36). Transcription of only two out of twenty genes close to *oriC* and a few outside a cluster of three *parS* sites was affected in a Δ*parB1* mutant of *Vibrio cholerae* (57). A similar limited regulation was also observed in *S. pneumoniae* (53) for the competence operon *com*, one of the operons localized in proximity of the *parS*s cluster.

*P. aeruginosa* is exceptional in this context, since deletion of the *parB* gene leads to global transcriptional changes affecting more than 1000 genes (58). It remains elusive how ParB of *P. aeruginosa* affects the expression of so many genes. Ten potential 16-bp palindromic *parS* sequences have been identified *in silico* in the genome of PAO1 strain of *P. aeruginosa*, numbered clockwise, with four of them (*parS1-parS4*) clustering next to *oriC* (31). *In vitro* studies have demonstrated that ParB binds to all ten *parS*s but with diverse affinity (59). The highest ParB binding affinity was found towards the perfect palindromic sequences *parS2* and *parS3*, slightly lower towards *parS1* and *parS4* with one mismatch, and even lower towards the remaining six *parS*s with two mismatches. *parS5* was the poorest binder of ParB *in vitro*. An analysis of our collection of single and multiple *parS* mutants has demonstrated that the presence of at least one high affinity site from the *parS1-parS4* cluster is necessary and sufficient for proper chromosome segregation (59). The remaining six sites were dispensable for chromosome segregation (59, 60).

Here we used chromatin (nucleoprotein) immunoprecipitation followed by next generation sequencing (ChIP-seq) to analyse the ParB-DNA interactions in *P. aeruginosa* wild type strain, a ParB-overproducing strain and several *parS* mutants: *parS*_null_ with all ten *parS* sites modified, *parS1-4* mutant with inactivated four *parS*s of the highest ParB affinity, and *parS2*^+^ with only one high affinity site left active in the background of the *parS*_null_ strain. This allowed us to identify a large number of additional, specific ParB binding sequences. Consequently, we propose that ParB, having hundreds of defined interaction sites in the genome, represents a new subfamily of Nucleoid-Associated Proteins (NAPs).

## MATERIALS AND METHODS

### Bacterial strains, growth conditions and plasmid manipulations

*Pseudomonas aeruginosa* PAO1161 (*leu*^-^ *r*^-^), a derivative of PAO1, was provided by dr. B.M. Holloway (Monash University, Clayton, Victoria, Australia). This strain lacks the inversion between the *rrn* operons present in the annotated PAO1 genome (61, 62). *P. aeruginosa* PAO1161 Rif^R^ (WT) strain (23), PAO1161 Rif^R^ *parB*_null_ (24), PAO1161 Rif^R^ *parS*mut15 with four modified *parS* sequences (here referred to as *parS1-4* mutant), PAO1161 Rif^R^ *parS*mut28 (here referred to as *parS2*^*+*^ mutant) with nine *parS* sequences modified but with unchanged *parS2*, and PAO1161 Rif^R^ *parS*_null_ with all ten *parS*s modified (59) were used in the analysis. A ParB-overproducing strain (designated ParB+++) was obtained by transformation of WT strain with pKGB9 (*araC-araBAD*p-*parB*). Chloramphenicol was added to 75 μg ml^-1^ (liquid cultures) or 150 μg ml^-1^ (solid media) to maintain the plasmid and 0.02% arabinose was added to induce the expression of *parB* (63).

### DNA pull-down assay

DNA pull-down analysis was carried out as previously (64) with modifications described in Supplementary Materials and Methods. Biotinylated DNA fragments containing *parS1* to *parS10* were obtained by PCR using appropriate pairs of primers (Supplementary Table S1). ParB binding to the DNA fragments was assessed by detecting ParB in the eluate using Western blotting with anti-ParB antibodies (24) and mass spectrometry analysis (Mass Spectrometry Laboratory, IBB PAS).

### Nucleoprotein immunoprecipitation (ChIP)

ChIP was performed as described previously (65) with modifications indicated in Supplementary Materials and Methods. Exponentially growing cultures in L broth at 37°C (OD_600_~0.5) were used. For each biological replicate, two independent 50 ml cultures of each strain were pooled together. The immunoprecipitation was performed with affinity purified rabbit polyclonal anti-ParB antibodies (31).

### ChIP-qPCR analysis

The immunoprecipitated DNA was used as a template in qPCR performed with Hot FIREPol EvaGreen qPCR Mix Plus (Solis Biodyne) and primers listed in Supplementary Table S1 in a Roche LightCycler 480. The efficiency of amplification for all primer pairs was between 1.95 and 2.05. The ChIP-qPCR results are presented as ChIP % recovery relative to the input DNA used in the procedure, calculated using the formula 100*2^(Ct_Input_ - 9.965 - Ct_ChIP_), where Ct – threshold values calculated using the 2^nd^ derivative method with LightCycler480 software ver 1.5.1.62. Before analysis the input samples were diluted 1000x. Each target sequence was analysed in samples from at least three biological replicates, each with three technical replicates. Primers amplifying a fragment of *proC* gene were used to assess non-specific (background) DNA recovery in the ChIP samples.

### ChIP-seq analysis

Library preparation and sequencing on Illumina HiSeq 4000 were performed at Genomed S.A. (Poland). Adapter sequences were removed using Cutadapt (66). Subsequent bioinformatic analyses were performed using the Galaxy platform (http://usegalaxy.org/) and R (67, 68). Reads were mapped to the *P. aeruginosa* PAO1 (NC_002516.2) genome using Bowtie2 (69). ParB peaks were identified by comparing combined replicates of IP samples (treatment) for each strain with combined negative control (*parB*_null_) samples using the callpeak function of MACS2 (70) with default options for paired-end BAM files, and 0.05 as FDR cut off. Visualisation of data was performed using DeepTools (71), Integrative Genomics Viewer ver. 2.3.91 (72) and Sushi (73). Coverage files, normalized to 1x sequencing depth (RPGC), were generated without binning and smoothing using the bamCoverage tool (71). For simplicity of presentation, the coverage values for individual nucleotides in biological replicates were averaged. ComputeMatrix and plotHeatmap components of deepTools (71) were used to prepare the heatmaps. Identification of the DNA motifs enriched in the ParB peaks was performed using MEME-ChIP (74). Differential binding analysis was performed using Diffbind (75).

## RESULTS

### ParB binding to *parS* sequences *in vitro* and *in vivo*

Bioinformatics analysis using the consensus *parS* sequence of *B. subtilis* (28) with allowance of two mismatches has led to the identification of ten putative ParB binding sites in the *P. aeruginosa* PAO1 genome (31). Our previous *in vitro* studies using electrophoretic mobility shift assay (EMSA) confirmed ParB binding to these sequences and demonstrated the highest affinity of purified His_6_-ParB to the *parS1-parS4* sites and the lowest to *parS5* (59). Since the EMSA was performed with short oligonucleotides and tagged ParB, here we used the DNA pull-down method to analyze the binding of WT ParB to different *parS* sequences in their natural context. Biotinylated DNA fragments (350-450 bp) encompassing *parS* sequences were attached to streptavidin-coupled magnetic beads, incubated with PAO1161 WT cell extracts and washed. Proteins bound to DNA were eluted and subjected to Western Blot analysis with anti-ParB antibodies as well as to mass spectrometry analysis. ParB was detected in the elution fractions in all samples containing DNA with any *parS* sequence with the exception of *parS5* and *parS8* (Figure 1A). These data indicate that in addition to the *parS1*-*parS4* sequences, required for accurate DNA segregation (59), ParB protein also binds predicted genomic *parS* sites: *parS6, parS7, parS9* and *parS10*.

**Figure 1.**
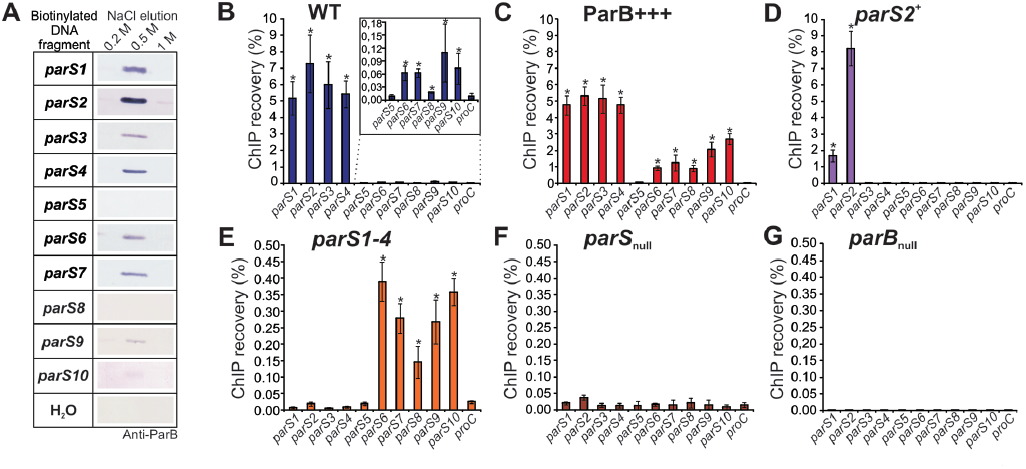
ParB binding to *parS* sites in *P. aeruginosa* genome. (**A**) Western blot analysis of elution fractions from DNA pull-down assays performed with DNA fragments containing *parS1 - parS10* sequences. Biotinylated DNA was coupled with magnetic beads, incubated with extracts of PAO1161 cells and washed. Proteins bound to the DNA were eluted with increasing concentration of NaCl (0.2-1 M). The negative control contained water instead of biotinylated DNA. Western blotting was performed using polyclonal anti-ParB antibodies. (**B-G**) qPCR analysis of DNA obtained by nucleoprotein immunoprecipitation using anti-ParB antibodies and extracts from exponentially growing cells of *P. aeruginosa* strains (**B**) WT, (**C**) ParB+++, (**D**) *parS2*^*+*^, (**E**) *parS1-4*, (**F**) *parS*_null_, and (**G**) *parB*_null_ (negative control). qPCR was performed using primers flanking the *parS* sequences and for the *proC* gene (background control). Data represent percentage of DNA recovered by ChIP relative to corresponding input samples and are shown as mean ±SD for at least three biological replicates analysed in three technical replicates. The significance of differences in recovery of *parS*s relative to the background control (*proC*) was evaluated by two-sided Student’s *t*-test assuming equal variance, with *p*<0.01 considered as significant (marked by asterix).

To verify the ParB binding to these *parS* sequences *in vivo*, chromatin immunoprecipitation (ChIP) with anti-ParB antibodies followed by quantitative PCR analysis was applied. Since a lack of ParB in *P. aeruginosa* PAO1161 *parB*_null_ mutant produced clear-cut effects under conditions of exponential growth in rich medium at 37°C, but not as severe as during growth in minimal medium (24, 76), such milder conditions were used for these experiments. The ChIP procedure was performed with PAO1161 (WT) strain, PAO1161 *parB*_null_ mutant (negative control), and PAO1161/pKGB9 (*araC-araBAD*p-*parB*) strain (ParB+++) with 5-fold increased ParB level which does not retard bacterial growth (63). Additionally, three *parS* mutants were included: *parS*_null_ with all ten *parS* sites mutated, *parS1*-*4* mutant with the four high affinity ParB sites inactivated, and *parS2*^+^ strain with nine *parS* sequences modified and only *parS2* left intact (59). Despite producing native amounts of ParB, both *parS1-4* and *parS*_null_ are defective in chromosome segregation to a similar extent as the *parB*_null_ mutant (24). The rationale behind using these strains was that it could help to detect ParB binding not only to the known *parSs* but also to other sites, by either increasing the cellular ParB level or by re-distribution of ParB from the main binding sites involved in chromosome segregation, as shown previously for a *parS*Δ*6* mutant of *B. subtilis* (36).

DNA recovered after ChIP as well as corresponding input DNA was used in quantitative real-time PCR with primers amplifying fragments encompassing *parS1* to *parS10* sequences and a fragment of *proC* gene as a background control (Supplementary Table S1). In WT samples strong ParB binding to *parS1* to *parS4* was confirmed as manifested by high ChIP % recovery (Figure 1B). Additionally, a statistically significant enrichment of sequences encompassing *parS6, parS7, parS8, parS9* and *parS10* but not *parS5* was found (Figure 1B, inset), indicating that under the growth conditions tested ParB did bind *in vivo* to all the *parS* sequences with the exception of *parS5.* Also in the ParB+++ strain ParB was bound to all the *parSs* but *parS5* (Figure 1C). Interestingly, while the regions encompassing *parS1* to *parS4* were recovered with a similar efficiency as in the WT, the remaining *parS* sites were significantly more enriched than in WT (Figure 1B and C). As expected, in the *parS2*^*+*^ strain, the region encompassing *parS2* showed the highest enrichment (Figure 1D), but also the DNA region around *parS1*, adjacent to *parS2*, was significantly enriched, suggesting that ParB bound to *parS2* spread to adjacent DNA sequences. In the *parS1-4* mutant the fragments containing *parS1* to *parS4* were not significantly enriched (Figure 1E) but those encompassing *parS6* to *parS10* were recovered with a higher efficiency than for the WT strain. Finally, no ParB binding to the altered *parS* sequences was found in the *parS*_null_ strain (Figure 1F), and for the *parB*_null_ strain (negative IP) the recovery was less than 0.01% for all the sequences analysed (Figure 1G).

Taken together, these results confirm that the *parS1* to *parS4* sequences are the major ParB binding sites in *P. aeruginosa* cells. The fact that the lower-affinity sites *parS6* to *parS10* were also enriched, and that this enrichment was higher in ParB-overexpressing cells and in cells lacking functional *parS1* - *parS4* sequences suggests that the ParB pool not bound by its major targets may more effectively associate with other sites of the bacterial chromosome.

### Genome-wide identification of ParB-enriched regions

To gain a global insight into the ParB interactions with the *P. aeruginosa* chromosome, chromatin immunoprecipitation was combined with next-generation sequencing (ChIP-seq). The anti-ParB immunoprecipitated DNA from two independent biological samples of each: WT, ParB+++, *parS2*^+^, *parS1-4, parS*_null_ and *parB*_null_ cells was sequenced and the reads were mapped to the *P. aeruginosa* PAO1 (NC_002516.2) genome. PlotFingerprint analysis indicated strongest local ChIP enrichment (a large part of the reads mapped to a few genomic bins) for WT strain and a more dispersed distribution of ParB-bound DNA regions in strains lacking functional *parS1* to *parS4* sequences and in the strain overproducing ParB (Supplementary Figure S1A). Some enrichment in specific regions of the genome was observed for the *parB*_null_ strain, which was likely due to cross-reactions of the polyclonal antibodies used.

A principal component analysis (PCA) on the mapped reads obtained for the six strains produced three clusters (Figure 2A). WT, ParB+++ and *parS2*^+^ formed one group, *parS1*-*4* and *parS*_null_ formed a second group, and the *parB*_null_ (negative control) strain was clearly separated from those two. Similar grouping was obtained by hierarchical clustering analysis based on correlation coefficients between different samples (Supplementary Figure S1B). Additionally, manual inspection of the distribution of the sequence reads along the reference genome showed that the reads from the five strains expressing ParB mapped roughly to the same regions which were different from the regions overrepresented in the *parB*_null_ strain (Supplementary Figure S1C). Overall, these analyses confirmed that the obtained reads were enriched in sequences specifically bound by ParB.

**Figure 2.**
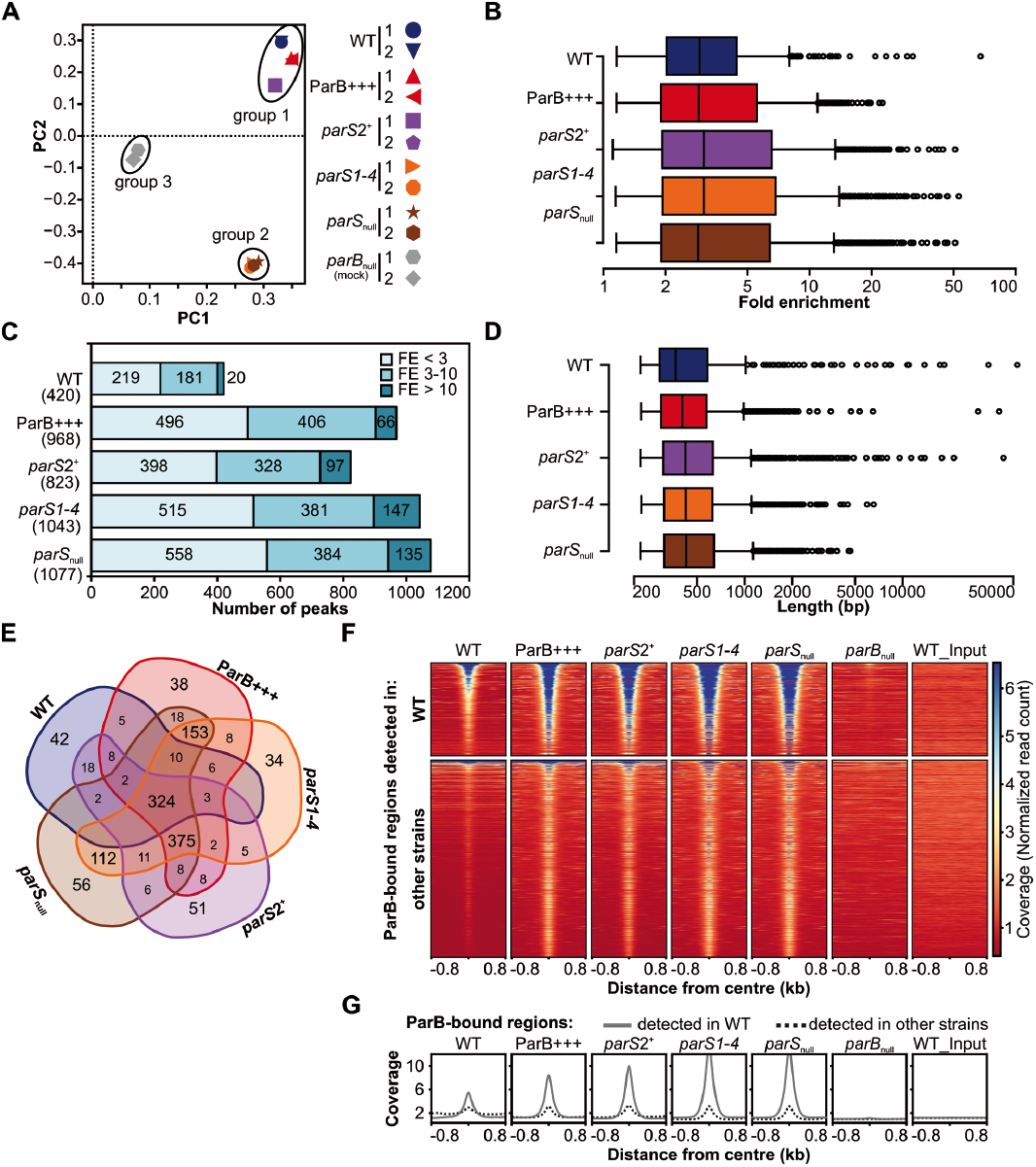
ChIP-seq analysis of ParB interactions with *P*. *aeruginosa* chromosome. Reads obtained by ChIP-seq for two biological replicates were mapped to the PAO1 genome using Bowtie2 (69) to generate binary alignment map (BAM) files, subsequently used in peak calling with MACS2. (**A**) Principal component analysis (PCA) of BAM files. (**B**) Distribution of fold enrichment (FE) values for detected peaks. (**C**) Number of ParB ChIP-seq peaks categorized according to their FE. (**D**) Distribution of lengths of ParB ChIP-seq peaks. (**E**) Compilation of ParB-bound regions in five strains. Two peaks in two strains were considered identical if their summits were less than 200 bp apart. The analysis yielded 1305 unique regions (Supplementary Table S7). (**F**) Coverage heatmap around centres of ParB-bound regions. The analysis was performed separately for 420 ParB-associated regions detected in WT strain (top) and 885 regions only detected in remaining four strains (bottom). Normalized read counts for each nucleotide were averaged for two biological replicates of each strain and coloured according to scale. Each line in the heatmap represents ±800 bp around the centre of the region. Rows were sorted in descending order of mean coverage value. The same regions analyzed in *parB*_null_ and WT input are included as controls. (**G**) Mean coverage for two groups of analysed regions in different strains.

To identify the sequences corresponding to the ParB binding sites we performed peak calling on the merged data for biological replicates, based on their high correlation (Supplementary Figure S1B and C). Using the FDR cut off value of 0.05, 420 specific peaks were found for the WT strain, 968 peaks for ParB+++, 823 for *parS2*^+^, 1043 for *parS1-4* and 1077 peaks for *parS*_null_ (Supplementary Tables S2-S6). The identified peaks in the WT displayed fold enrichment (FE) between 1.15 and 67.5, with a median of 2.9 (Figure 2B). A similar median FE (3-3.1) was observed for all the other strains analyzed (Figure 2B). Remarkably, not only the number of peaks but also the number of peaks with FE >10 was substantially higher in the ParB+++ strain as well as in the strains with the high-affinity *parS* sites impaired (Figure 2C), indicating feasibility of the strategy applied for identification of additional ParB binding sites.

The lengths of the sequences encompassed by the peaks varied greatly between 0.2 kb and about 50 kb in WT, ParB+++ and *parS2*^*+*^ samples (Figure 2D). In *parS1-4* and *parS*_null_ the broadest peaks reached only 8 and 5 kb, respectively. Nevertheless, the median values of the peak width did not differ substantially among the strains (Figure 2D).

The peaks obtained for the five strains were matched based on the proximity of peak summits (Supplementary Table S7). Approximately 90% of the corresponding peaks in various strains had their summits less than 20 bp apart. Overall, the analysis yielded 1305 groups of peaks representing regions with increased ParB occupancy (Supplementary Table S7, hereafter referred to as ParB-bound regions). Of those, 324 were common to all five strains, representing a high-confidence set of ParB-bound regions in the *P. aeruginosa* genome (Figure 2E, Supplementary Table S7). Three other most numerous pools consisted of: 375 sites identified in four strains (ParB+++ and the three *parS* mutants), 153 in three strains (ParB+++, *parS1-4* and *parS*_null_), and 112 sites common to *parS1-4* and *parS*_null_. Less than 10% of the peaks were unique to any single strain.

ParB-bound regions generally displayed an increased read coverage in the central parts (Figure 2F and 2G). The 420 regions identified in WT strain (Figure 2F, top) were also occupied by ParB in all the other strains, with the exception of those harbouring *parS* sequences that were inactivated in the respective *parS* mutants. Importantly, no increased coverage was visible for *parB*_null_ or WT_input. Similar inspection of ChIP-seq data for remaining 885 ParB-bound regions detected only in ParB+++, *parS2*^+^, *parS1-4* or *parS*_null_ revealed that most of these regions also displayed a slight increase of coverage in WT (Figure 2F, bottom and 2G), suggesting that ParB could also bind to these sites in WT cells. Overall, this data indicated the existence of more than 1000 specific ParB-bound regions in the *P. aeruginosa* genome. The high reproducibility of these regions in different *P. aeruginosa* strains implies robust and specific binding of ParB to genomic DNA.

### Inspection of ParB ChIP-seq peaks in various strains

Distribution of the ParB-enriched regions in various strains was visualized on the *P. aeruginosa* PAO1 genome (Figure 3A). For clarity, only peaks with FE >3 were considered. The ParB-bound regions were present across the entire genome, but the highest peaks (marked with dots) were almost exclusively found in the genome half containing *oriC*. Notably, the broadest peaks (>5 kb), almost exclusively found in only three strains, WT, ParB+++ and *parS2*^+^, were tightly clustered in less than ca. 8% of the genome around *oriC* (triangles in Figure 3A). In the *parS1-4* mutant only two peaks broader than 5 kb were detected. Interestingly, these peaks encompass *parS6* and *parS10*.

**Figure 3.**
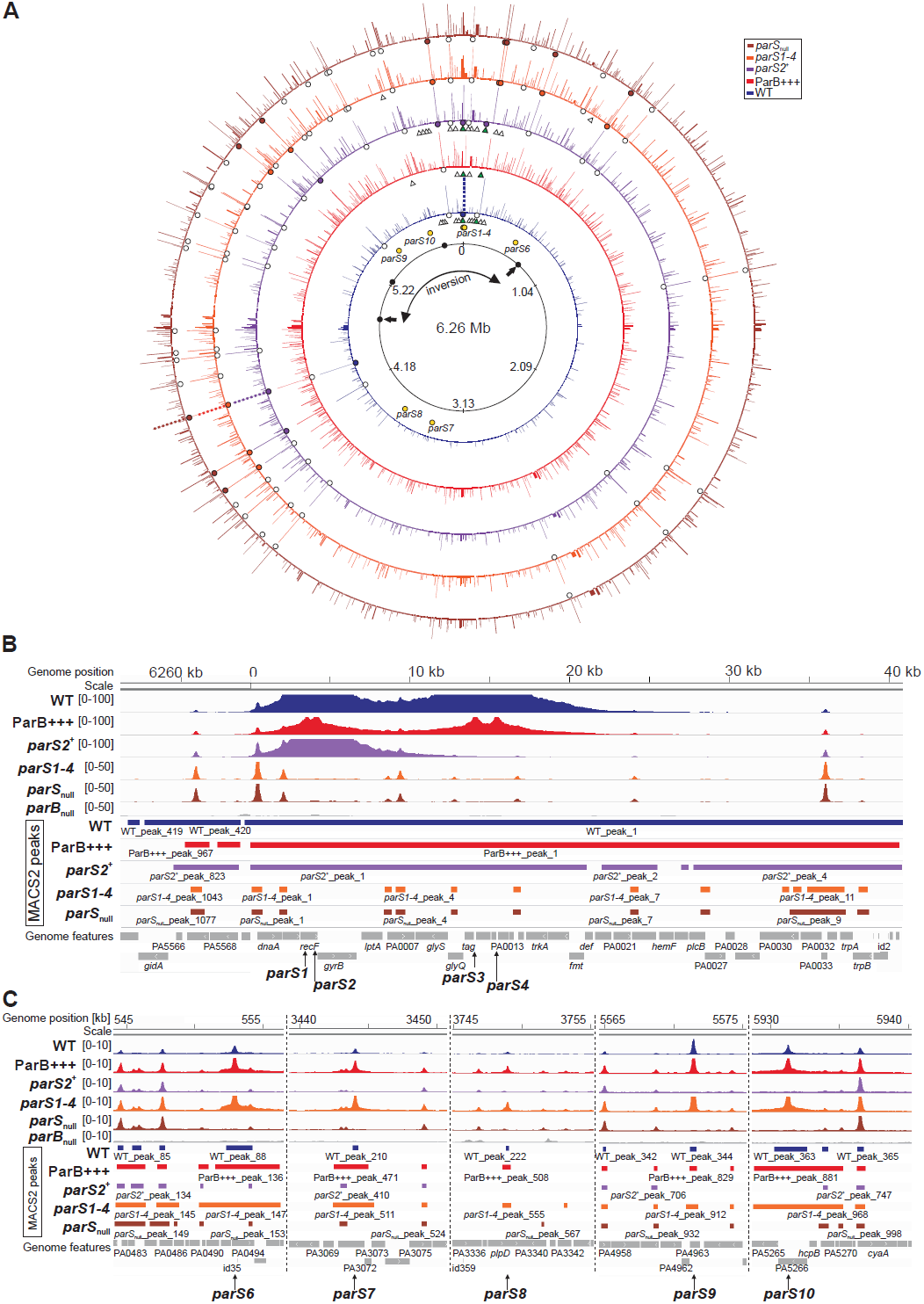
Localization and characteristics of ParB ChIP-seq peaks. (**A**) Distribution of peaks detected in five analysed strains. For simplicity, only peaks with fold enrichment >3 are shown and all peaks with FE ≥30 are shown at the same height. White dots below peaks indicate those with 20≤ FE ≤30, coloured dots those with 30< FE <50, dotted lines mark the highest peaks with FE ≥50. White triangles mark peaks broader than 5 kb, green those >20 kb. *parS* sequences are indicated with yellow dots. Genomic coordinates are according to the PAO1 genome. The *rrnA - rrnD* gene clusters are marked with black dots, black arrows indicate *rrnA* and *rrnB* loci sites where inversion occurred in PAO1 (61, 62) but not in PAO1161 strain. (**B**) Coverage of the genome regions encompassing *parS1* - *parS4* cluster and (**C**) *parS6* - *parS10* sequences in each strain. The histograms, show normalized read counts (averaged for two biological replicates) for corresponding positions of the PAO1 genome. Ranges of peaks and annotated genomic features are indicated below.

The 20 peaks with FE >10 identified in the WT (Figure 2C) were analyzed in detail in the four other strains (Table 1). The cluster of four high affinity *parS* sites (*parS1* - *parS4*) in the WT strain produced an extended peak of 53 kb with the maximum FE of 67 (Figure 3B). The corresponding 50-kb peak was only 20-fold enriched in ParB+++ strain. In the *parS2*^+^ mutant a peak with the summit at *parS2* encompassed 21 kb and had FE of 33. No such broad peaks were detected in the *parS1-4* or *parS*_null_ strains but the absence of ParB binding to the *parS1*-*parS4* cluster in these mutants unmasked additional 15 or 16 peaks, respectively, one of them upstream of *dnaA* with a FE ca. 19. The role of the ParB binding in the expression of *dnaA* operon requires further studies.

**Table 1.**
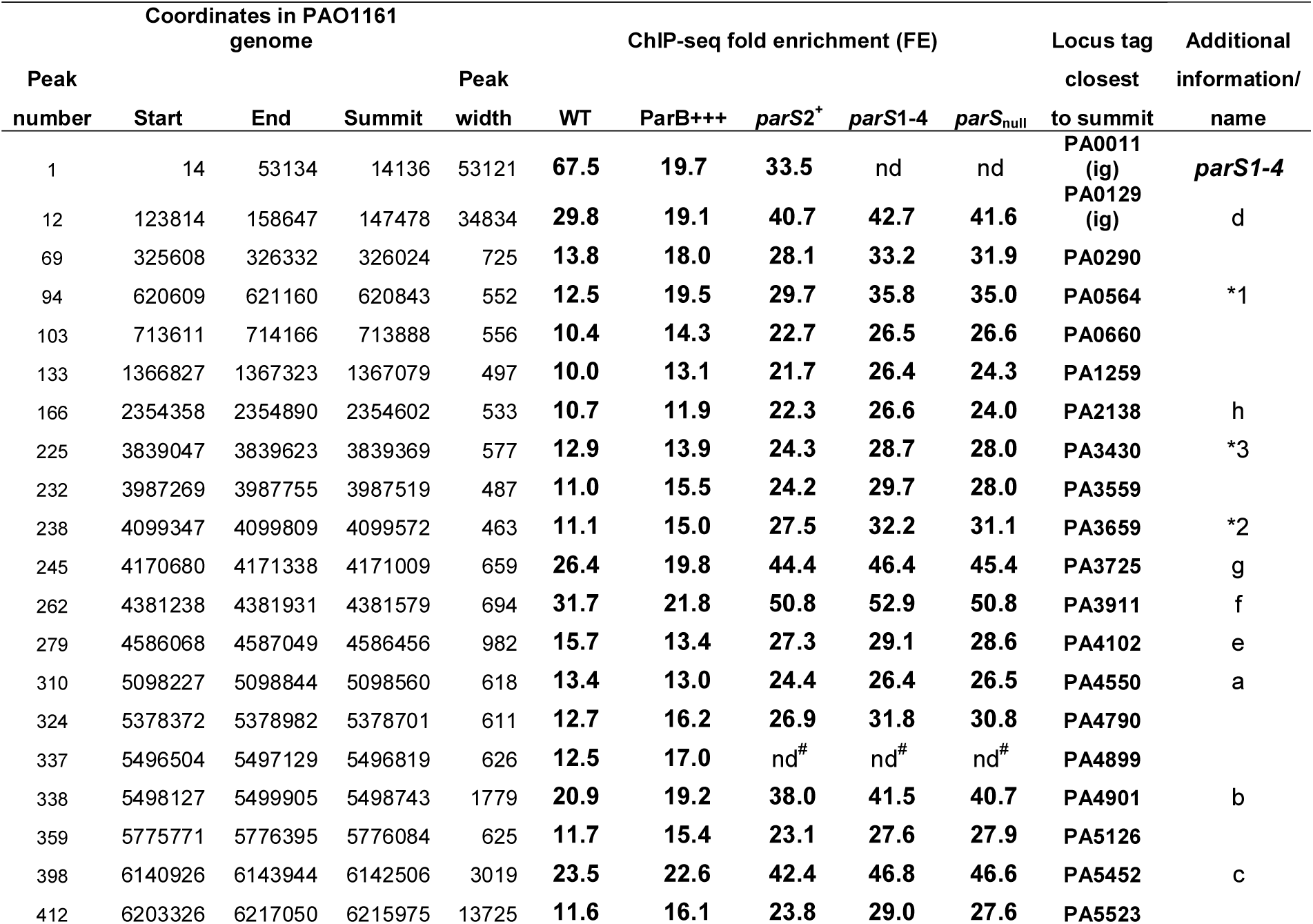
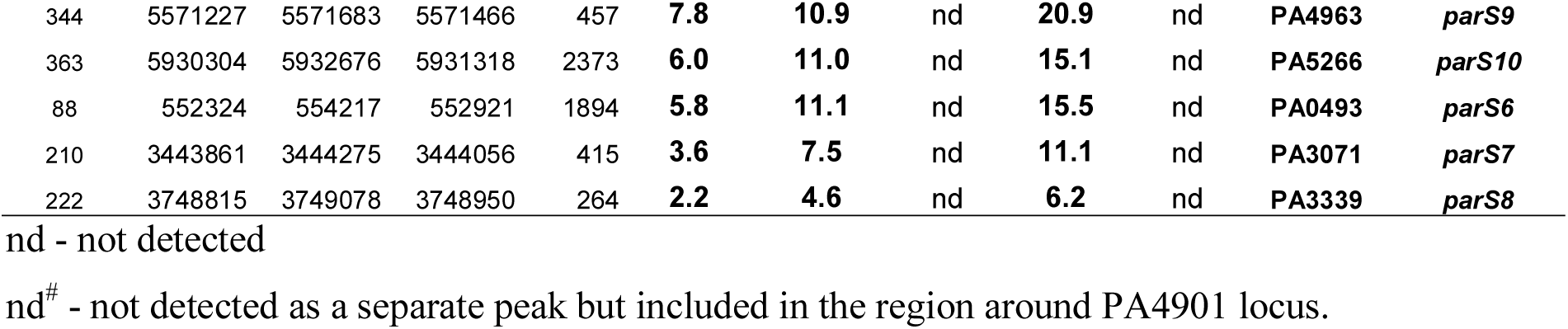
Selected ParB-bound regions in *P. aeruginosa*. Shown are parameters of ParB ChIP-seq peaks with fold enrichment higher than 10 as determined by MACS2 callpeak analysis of ChIP-seq data for *P. aeruginosa* PAO1161 (WT) strain using *parB*_null_ strain as a control. Corresponding peaks in other strains were identified by peak matching (Supplementary Table S7). Data for peaks containing *parS6 - parS10* sequences are also included. Last column provides names of *parS* sequences and designations of the ParB binding sites identified before (60).

The remaining 19 highest peaks detected in WT strain were also present in the four other strains and had the FE >10. Interestingly, in the three *parS* mutants the accumulation of ParB at these 19 loci was much higher than in WT or ParB+++ (Table 1). Notably, none of these 19 ParB-associated peaks corresponded to known *parS* sequences. Nevertheless, the *parS6* to *parS10* sites (with the exception of *parS5*) were ParB-bound in three strains, with 3-to 4-fold higher ParB occupancy in ParB+++ and *parS1-4* strains compared to WT (Table 1; Figure 3C).

The width of the *parS*-encompassing peaks varied for different *parS*s and also depended on the relative availability of ParB. The *parS6* and *parS10* containing peaks encompassed ca. 2 kb each in WT, but were ca. 6 kb wide when an excess of ParB was available, i.e., in ParB+++ and *parS1-4*. ParB bound to *parS7, parS8* and *parS9* occupied ~400 bp in WT and under conditions of ParB overabundance the ParB-bound region extended significantly (~3 kb) only for *parS7*. These data suggest that ParB bound to a *parS* site can spread for variable distances depending on the individual *parS* sequence and the availability of ParB.

Overall, the ChIP-seq analysis demonstrates that *in vivo* ParB binds not only to the *parS* sequences identified previously, with the exception of *parS5* (Figure 3B and C; Table 1), but also to additional regions across the entire genome creating a specific, highly reproducible pattern of distribution (Figure 3A).

### DNA motif other than palindromic *parS* is bound by ParB in *P. aeruginosa* genome

To check if the genome-wide ParB binding involved a defined DNA motif, we performed a search of DNA sequences flanking the summits (±75 bp) of the peaks using MEME ChIP (74). Remarkably, a similar 7-bp DNA sequence logo was identified in data from all five strains (Figure 4A). The GtTcCAc motif showed conservation of positions 1, 3, 5 and 6 whereas at the three other positions some freedom (T→C/A at position 2, C→T at positions 4 and 7) was allowed (Figure 4A). Notably, this motif resembles GTTCCAC sequence, which is a conserved part of the 8-bp inverted arms of the four major *parSs* critical for chromosome segregation, *parS1* - *parS4* (Figure 4B), and is also present in one arm of the remaining *parS* sequences, with the exception of *parS5*, the only *parS* to which ParB binding was not confirmed (Figure 1). In *parS6* - *parS10* the other arms carry substitutions at positions 1, 3 and/or 6, the positions strictly conserved in the deduced motif (Figure 4B), suggesting that these parts of *parS6*- *parS10* might not be involved in initial ParB binding.

**Figure 4.**
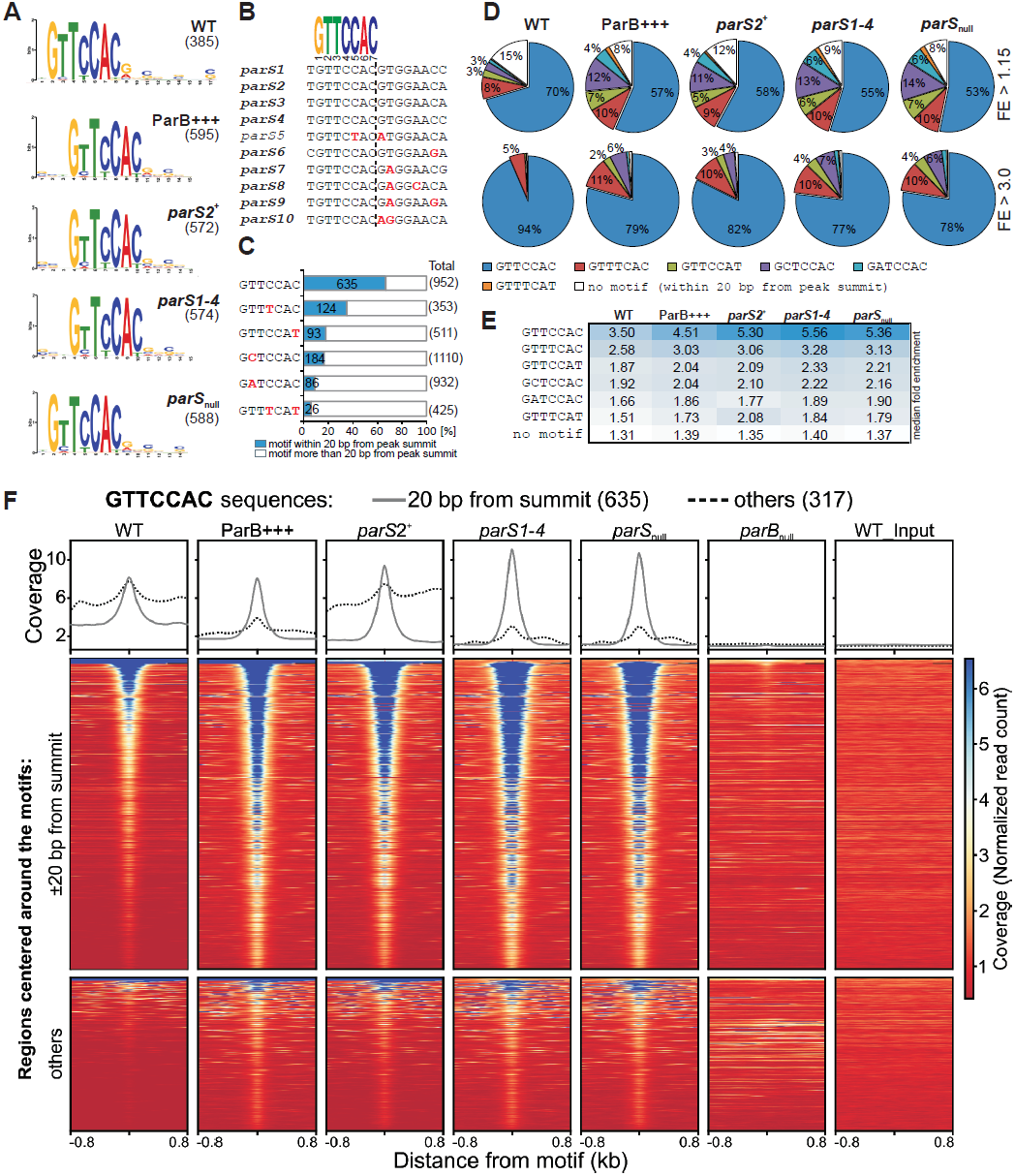
Recurrent sequence motifs in ParB-bound regions of *P. aeruginosa* genome. (**A**) Sequence logos of DNA motifs enriched in genome sequences ±75 bp from summits of ParB peaks identified in indicated strains. Motif search was performed using MEME-ChIP. The number on the right indicates the total amount of sequences used to build the logo. (**B**) Alignment of *parS* sequences from the *P. aeruginosa* PAO1 genome. Logo represents core sequence conserved in both arms of *parS1* - *parS4* palindromes and in one arm of the remaining *parS* sequences, with the exception of *parS5*. Mismatches relative to GTTCCAC are indicated in red. (**C**) GTTCCAC sequence variants enriched around summits of ParB ChIP-seq peaks. The number of the sequences found within ±20 bp of any peak summit relative to the total number of copies in the PAO1 genome is shown. Only motifs enriched more than 4 times are presented. Data for remaining variants are included in Supplementary Table S8. Mismatches relative to GTTCCAC are indicated in red. (**D**) The percentage of peaks containing the indicated heptanucleotide motifs as closest to the summit. The analysis was performed for all peaks (upper panel) and for peaks with FE >3 (bottom panel). Category “no motif” includes peaks with none of the analysed variants present ±20 bp from summit. (**E**) Median fold enrichment of ParB ChIP-seq peaks carrying indicated variant of the heptanucleotide motif as closest to the summit in different strains. (**F**) Read coverage around GTTCCAC motifs in the PAO1 genome in ChIP samples from different strains. Each line in the heatmap represents normalized read counts for each nucleotide of ±800 bp region around the GTTCCAC motif sorted in descending order of mean coverage value, averaged for two biological replicates and coloured according to scale. Upper part corresponds to fragments with GTTCCAC motif identified within 20 bp from ParB peaks summit, bottom part to fragments with remaining GTTCCAC motifs. The same regions analyzed in *parB*_null_ and WT input are included as controls. Plots above the heatmaps indicate mean coverage score separately for the two groups of GTTCCAC motifs.

To identify the sequence variants which could be recognized by ParB, a search of DNA motifs in the PAO1 genome was performed using the GTTCCAC sequence with the allowance of two mismatches. The total pool of such variants was then limited to the 183 ones detected at least once within 20 bp from any peak summit (Supplementary Table S8). Overall, the sequences in proximity of peak summits analysed here represent 1.3% of the genome. Remarkably, six variants: GTTCCAC, GTTTCAC, GTTCCAT, GCTCCAC, GATCCAC and GTTTCAT showed at least 4-fold enrichment in sequences ±20 bp of peak summits relative to the total genome (Figure 4C, Supplementary Table S8). Among them, GTTCCAC was most overrepresented, since 66.7% from 952 sequences present in the PAO1 genome was found close to peak summits. The second motif most enriched in proximity of summits was GTTTCAC (35.1%).

Analysis of the peaks in all five strains revealed that ≥85% contained at least one GTTCCAC, GTTTCAC, GTTCCAT, GCTCCAC, GATCCAC or GTTTCAT motif within 20 bp from the summit (Figure 4D). GTTCCAC was the most frequently identified variant localized closest to the summit (Figure 4D, upper panel), followed by GTTTCAC. Remarkably, when the analysis was limited to the peaks with FE >3 these two variants were identified in 94% and 5% of WT peaks, respectively (Figure 4D, bottom panel). Similar trend was observed for other strains. Strikingly, peaks without any of the six analysed motifs ±20 bp from summit represented less than 1% of the peaks with FE >3 (Figure 4D, bottom panel).

A complementary analysis of the fold enrichment of the peaks containing indicated variant as the closest one to the summit gave the highest median FE for peaks with GTTCCAC, followed by those with GTTTCAC (Figure 4E). The median fold enrichment for the peaks without motif ±20 bp from summit was less than 1.4 suggesting that these peaks might represent low affinity binding sites (Figure 4E).

To further confirm that the genome fragments carrying the six indicated sequences were enriched by precipitation with anti-ParB antibodies we analysed normalized read coverage around each motif in the genome, separately for those found ±20 bp from the summits and for the remaining ones. As expected, the mean normalized read count around GTTCCAC present ±20 bp from a peak summit revealed a strong central increase in all ParB-expressing strains, but not for *parB*_null_ (negative control) or WT_input DNA (Figure 4F, upper part of the heatmap). Similar analysis for remaining GTTCCAC motifs also revealed an increase in read coverage for sequences in proximity of most of these motifs (Figure 4F, bottom part of the heatmap). The observations were confirmed by quantification of the coverage data for both groups (Figure 4F, plots). Similar results were obtained for GTTTCAC, GCTCCAC, GTTCCAT, GCTCCAC and GTTTCAT motifs (Supplementary Figure S2). Overall, the high enrichment of six sequence variants, related to one arm of *parS*, in proximity of ChIP-seq peak summits strongly suggests that these are true targets for ParB binding.

### ParB bound to *parS1-4* cluster provides a platform for extended ParB - DNA interactions beyond lateral spreading

The presence of the 7-bp motif in ParB-bound regions positively correlated with the number of strains in which the region was identified (Figure 5A). Out of the 324 ParB-associated regions common for all strains, 321 (99.1%) contained the motif in comparison with only 41% found in any single strain. Overall, the six motif variants were identified ±20 bp from summits of peaks in 1097 out of 1305 ParB-bound regions (Figure 5B) and in the next 59 ParB-bound regions the motifs were found more than 20 bp away from summits (Figure 5B). Only 149 regions lacked the analysed motifs. Of these, 70% were found only in a single strain (Supplementary Table S7) and corresponded to low-enriched regions. Notably, most of the motif-less peaks were detected in the WT, ParB+++ and/or *parS2*^+^ strains, i.e., the strains carrying at least one *parS* from the *parS1-4* cluster (Figure 5B). Almost all these peaks were in vicinity of *oriC* (Figure 5C). This suggests that formation of a large nucleoprotein complex after ParB binding to the high affinity site(s) (Figure 3B) caused certain DNA regions fairly distant from the *parS1*-*4* cluster and lacking the 7-bp motif to be recovered in the anti-ParB immunoprecipitate.

**Figure 5.**
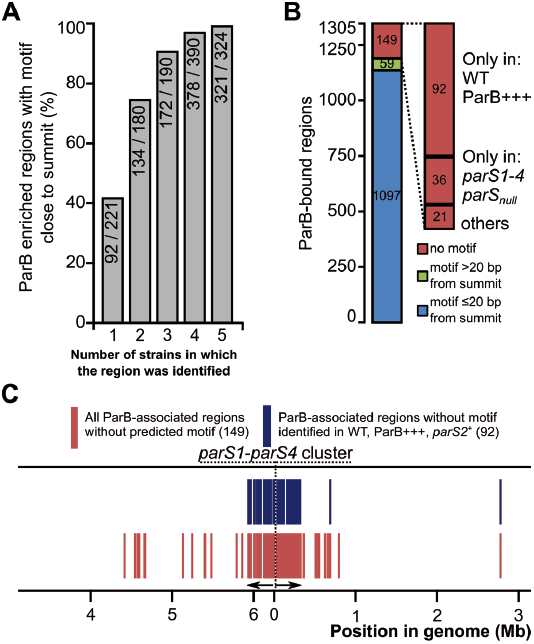
Heptanucleotide motif-independent ParB binding to DNA. (**A**) Fraction of ParB-associated regions carrying six analysed variants of heptanucleotide motif ±20 bp from peak summit as a function of number of strains in which the region was detected (Supplementary Table S7). (**B**) Distribution of heptanucleotide motifs (as above) in ParB-associated regions. The regions are divided into three categories: with a motif ±20 bp from summit, with a motif within the region but further away than 20 bp from summit, and with no motif. The motif-less regions were also categorized according to the strains they were detected in. (**C**) Localization of ParB enriched regions lacking a heptanucleotide motif in the *P*. *aeruginosa* PAO1 chromosome.

To get an insight into the postulated *parS1-parS4* dependent enrichment of specific DNA sequences a differential binding analysis was performed using Diffbind (75). ChIP-seq data for strain carrying at least one *parS* of the *parS1-parS4* cluster (WT, ParB+++ and *parS2*^+^) was compared with data obtained for strain lacking such *parS* site (*parS1*-*4* or *parS*_null_). The analysis was not limited to the ChIP-seq peaks, but instead non-overlapping 200-bp windows covering the whole genome were used. The WT *versus parS1-4* comparison revealed that the majority of bins showing statistically significant difference between the strains, displayed decreased coverage in WT (Figure 6A). This was in agreement with the previous observation that the ChIP-seq peaks in the *parS* mutants showed higher fold enrichment than the corresponding peaks in the WT (Table 1). Remarkably, some bins showed an increased coverage in WT relative to *parS1-4* and these were confined to a 400-kb chromosome region encompassing *oriC* (Figure 6A). The differential region including *oriC* displayed the highest differences from 10 kb upstream of *parS1* to 20 kb downstream of *parS4* (Figure 6B). If ParB was spreading, starting from *parS1-parS4*, the negative correlation between the magnitude of the observed difference between the strains and the distance from the *parS1*-*parS4* cluster should be observed. Remarkably, no such correlation was observed in the distant regions (e.g. 115 kb to 160 kb) as multiple bins in these regions showed higher coverage than bins located relatively closer to *parS1*-*parS4* (Figure 6A). Similar results were obtained when WT was compared with *parS*_null_ as well as when *parS2*^+^ or ParB+++ was compared with *parS1-4* (Supplementary Figure S3). Significantly higher coverage of sequences from the 115 kb to 150 kb region in WT *versus parS1-4* comparsion as well as presence of 20 kb gap separating this region from peak encompassing *parS1*-*parS4* were also observed when WT_input was used as a background control in Diffbind analysis, indicating that the observed distribution is not an artefact caused by an elevated number of reads mapping to this part of the genome in *parB*_null_ data, used as a background control in all previous analyses.

**Figure 6.**
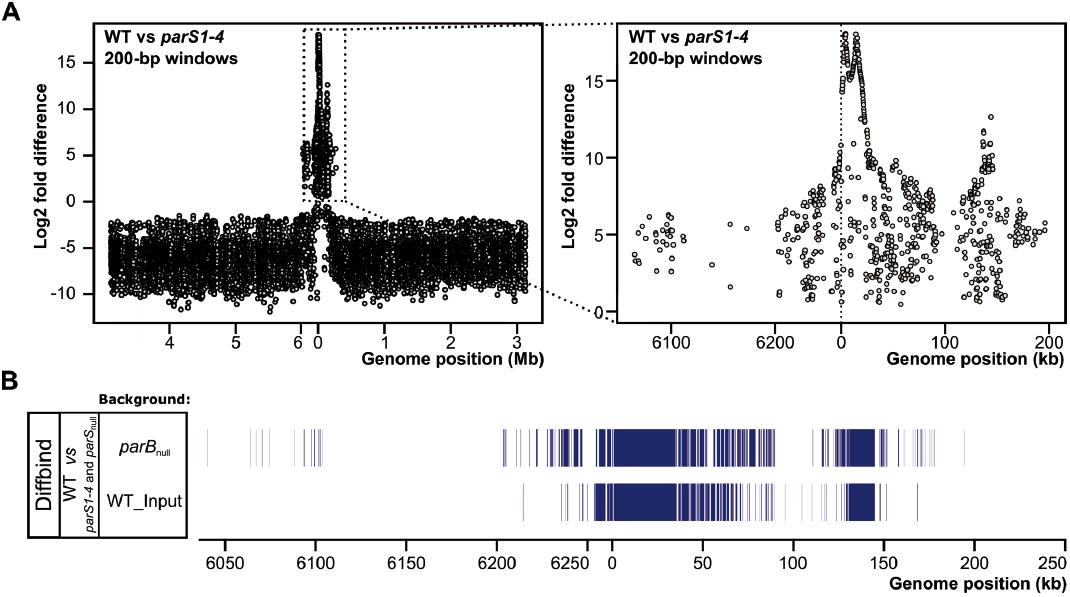
*parS1*-*parS4* dependent ParB distribution in *P. aeruginosa* genome. (**A**) Comparison of ParB binding to DNA in WT strain *versus parS1*-*4* strain (lacking functional *parS1* - *parS4* cluster). Differential binding analysis was performed in non-overlapping 200-bp windows covering the whole genome using Diffbind. Only bins showing statistically significant difference of coverage (FDR of 0.05) between WT and *parS1*-*4* are shown. On the right, blow up of the *oriC* region. Only bins showing higher number of reads in WT than in *parS1*-*4* are shown. (**B**) Distribution of *parS1*-*parS4* dependent ParB ChIP enriched sequences in proximity of *oriC* (±250 kb). Diffbind analysis was performed for WT *versus parS1*-*4* and *parS*_null_ ChIP-seq data using 200-bp windows. Subsequently the analysis was performed with the 200-bp windows shifted by 100 bp. Data represent windows (or their halves) showing the statistically significantly (FDR of 0.05) higher coverage with reads in WTstrain relative to *parS1*-*4* and *parS*_*null*_ for both sets of genomic windows using *parB*_null_ or WT_Input data as a background control.

Overall, the data indicate that ParB binding to *parS1*-*parS4* is required for an enrichment of additional specific chromosome regions around *oriC* but also quite distant from it. The fact that these regions are discontinuous suggests that the enrichment cannot be a consequence of lateral spreading of ParB along the DNA adjacent to the *parS1-4* cluster but supports the hypothesis of interactions at a distance, involving for example bridging/looping or caging of DNA.

## DISCUSSION

Bacterial genomes are highly compact structures whose distribution within cell is tightly controlled spatially and temporally (77, 78). Chromosome segregation must be coordinated with DNA replication and cell division to assure delivery of the full genome copy to progeny cells. It has been demonstrated that ParA, ParB and *parS* sequences are engaged in genome partitioning in many bacteria (9, 12–23, 25, 31). Additionally, they have been shown to be involved, in a species-specific manner, in the regulation of DNA replication initiation by DnaA (14, 15), in the formation of a loading platform for SMC on newly replicated DNA to assure its proper condensation (46, 47), in positioning of *oriC* domains at defined cell locations (9, 12, 24, 25, 43, 44, 79), coordination of replication and cell division (18, 39, 80) but also in regulation of gene expression (53, 57, 63, 80).

Our earlier studies have confirmed the partitioning role of ParA and ParB proteins in *P. aeruginosa*, identified ten *parS* sequences bound by ParB with different affinities and showed that any one of the four *parS* sites closest to *oriC* – *parS1*-*parS4* is required and sufficient for accurate chromosome segregation (31, 59). Remarkably, expression of over 1000 genes was affected by a lack of ParB (58), suggesting that ParB, directly or indirectly, plays a role in global gene regulation. Other plausible explanation of that effect invoked ParB interactions with sites other than *parS*s in the genome and/or an involvement of ParB partners in the regulation. However, a recent study on ParB in *P. aeruginosa* (60) claimed that *in vivo* ParB bound to the *parS1*-*parS4* sites but not to other *parS*s. Interestingly, eleven other regions were identified as ParB-bound but they were not thoroughly analysed. To solve this apparent discrepancy regarding the ParB - DNA binding *in vivo* we performed ChIP-seq using a WT strain, three *parS* mutants as well as a ParB-overproducing strain, with *parB*_null_ serving as a negative control. Unlike in the aforementioned study (60), native untagged ParB and polyclonal anti-ParB antibodies were used rather than a ParB-FLAG chimera. Additionally, we grew the cells in rich medium, since the mutants studied displayed much milder growth defects in such medium relative to minimal medium (24). During data analysis we did not apply strict fold-enrichment cut off, which allowed us to identify high number of ChIP-seq peaks. Importantly, more than 90% of the peaks were also identified in at least one other strain (Figure 3E) indicating that, despite relatively low FE, these peaks represent bona fide ParB-binding regions.

The ChIP-seq analysis showed unequivocally that *in vivo* ParB is bound to all the palindromic *parS* sequences identified previously, except *parS5*, with the highest degree of ParB association in proximity of *oriC* coinciding with the *parS1*-*parS4* sites (31, 59). Introducing mutations in the *parS* palindromes blocked the ParB binding which confirmed that the indicated *parS* are bound by ParB *in vivo*. Remarkably, our data indicate that ParB occupies 420 regions in WT strain, covering almost 7% of the genome and app. 1000 regions under conditions of ParB abundance.

Using stringent enrichment criterium of FE >10 there were 20 high-affinity regions identified in WT, including the *parS1-4* region and all eleven “secondary ParB binding sites” detected previously (60) in *P. aeruginosa* strains grown on minimal medium (Table 1). This correspondence of two sets of results indicates a high degree of specificity and conservation of ParB binding to the genome under various growth conditions. However, the drastically different experimental setups and approaches used in data analysis prevent conclusions about growth conditions dependent ParB binding to DNA.

The extent of ParB spreading has been analysed in diverse chromosomal systems. Most data suggested the spreading for up to 10 kb from a *parS* site in natural conditions (36, 53, 57, 80). Recently, ParB from *C. crescentus* has been shown to spread no more than 2 kb from a single *parS* site, while the extension of the ParB-associated region in the vicinity of *oriC* to 10 kb involved ParB binding to a cluster of four *parS* sequences located in a 5-kb region near *oriC* (50). The *parS1-parS4* cluster located within 12 kb of *oriC* in *P. aeruginosa* seems to facilitate even farther ParB - DNA interactions since the most conspicuous region of ParB spreading encompassed more than 50 kb in the WT strain. A comparison of the distribution of the ParB-associated DNA in the *oriC* region between the WT and *parS1-4* (or *parS*_null_) revealed a discontinuous stretch around the *oriC* whose occupation was strictly dependent on the presence of *parS1*-*parS4* (Figure 6). The occurrence of substantial ParB-free gap in this stretch as well as lack of correlation between magnitude of ParB enrichment and the distance from *parS1-4* sites, contradicted lateral spreading but was consistent with bridging and looping at long distances between ParB complexes shaping the ori domain (37, 51, 52, 81, 82).

Notably, we show that ParB peaks contain a motif GtTcCAc corresponding to seven out of eight nucleotides from a single arm of the canonical *parS* palindrome (Figure 4A). A systematic analysis of the motif variants with the allowance of two nucleotide substitutions showed that GTTCCAC, GTTTCAC, GCTCCAC, GTTCCAT, GCTCCAC and GTTTCAT motifs were significantly more often found in proximity of ParB ChIP-seq peak summits. In any given strain more than 85% of all peaks and 99% of peaks with FE >3 carried one of the six sequence variants in proximity of peak summit. Among these six variants, GTTCCAC was not only the most frequently occurring one but was also found closest to the summits of peaks with the highest FE. In the most consistent peaks, those detected in four or five strains, 97% and 99%, respectively, contained one or more of the six motif variants within 20 bp from the summit (Figure 5A). Remarkably, inspection of read coverage around all indicated motifs in PAO1 genome showed an increased coverage around these sequences indicating that these motifs could be targeted by ParB *in vivo*. Recent reports indicate that some ParB proteins may not strictly require the symmetrical *parS* structure for DNA binding. ParB2 of *Vibrio cholerae* was shown to bind a non centromeric site containing a half-*parS* sequence (80). Additionally, in multipartite genome bacterium *Burkholderia cenocepacia* ParBc1, ParBc3 and ParBpBC display enhanced affinity for the corresponding half-*parS* relative to the background control (83). This data suggests that ParB binding to one arm of *parS* may not be restricted to *P*. *aeruginosa* ParB. The biological significance of ParB binding to half-*parS* remains elusive. We speculate that half-*parS* sequences scattered across the genome may play a role in concentration of ParB molecules on the nucleoid. ParB interaction with multiple sites along with the interactions between ParB molecules may provide a scaffold important for chromosome compaction, proper DNA topology, possibly playing a role in modulation of gene expression. Overall, *P*. *aeruginosa* ParB may be classified as a nucleoid associated protein (NAP).

When the availability of ParB in the cell is limited, as is the case in the native situation, the protein must discriminate between its binding sites. Occupation of certain sites may be determined by the degree of affinity, interactions with cellular partners (e.g., ParA), local ParB availability, on-going DNA transactions (e.g., transcription and replication), presence of other factors interacting with DNA, DNA conformation, physiological state of the cell, or growth conditions. An increased ParB availability in the cells enabled identification of ParB binding to additional 885 loci not identified in the WT strain. Importantly, most of these loci show a slight increase in read coverage in WT ChIP-seq data suggesting the possibility of ParB binding to these sites in WT cells and not only in mutants used in this study (Figure 2F). Concomitantly, most of the peaks common for WT and mutant cells showed higher fold enrichment in the four strains with increased ParB availability. In a similar study in *B. subtilis* deletion of six out of ten *parS* sites resulted in re-distribution of Spo0J (ParB) towards the low-affinity *parS* sites left in the mutant strain (36), whereas in *P. aeruginosa* inactivation of the high affinity *parS* sites led to the re-distribution of ParB not only to the remaining *parS*s but also to hundreds of additional ParB binding sites.

Our earlier transcriptomic analysis of the *parB*_null_ mutant versus WT indicated a global regulatory role of ParB in gene expression in *P. aeruginosa* (58). A large fraction of the 1166 genes with altered expression seemed to be involved in the stress response due to the disturbances in chromosome segregation (84). A complementary study with a strain showing 5-fold overproduction of ParB (ParB+++), a ParB excess which did not lead to visible changes in growth rate or genome segregation, revealed altered expression of 211 loci (63). The present ChIP-seq data suggested that ParB-DNA binding relies on a presence of specific sequence(s). A comparison of the distribution of analysed variants with the transcriptional organization of the genome revealed that in PAO1 these sequences are mainly intragenic (GTTCCAC – 92%, GTTTCAC – 76%, GCTCCAC – 89%, GTTCCAT - 94%, GCTCCAC – 93% and GTTTCAT – 84%). The loci, which expression is affected by ParB deficiency are putatively expressed from 853 promoters. Only 113 of these promoters contain one of the six indicated sequences. Similarly, the 211 loci affected by a slight ParB overproduction expression are expressed from 130 promoters but only 28 of these promoters contain one of the indicated heptanucleotide motif variants. Whereas it was demonstrated that ParB binding to palindromic *parS3* and *parS4* was responsible for the repression of *orf011*p and *orf013*p, respectively (63), low correlation between the ParB dependent change in the expression and presence of half-*parS* sequences in the promoters of other genes suggest that ParB binding to half-*parS* motifs may not be the sole factor in regulation of the activity of these promoters. Further studies are required to gain insight into the mechanism of the postulated ParB-dependent transcription regulation.

Summarizing, the most prominent features of the *P. aeruginosa* partitioning protein ParB are: i/ *parS* sequence-specific DNA binding dedicated primarily to the specific task of *oriC* separation and chromosome segregation; ii/ specific binding to numerous heptanucleotide sites throughout the genome; iii/ acting at a distance, probably by a combination of spreading and bridging/ looping/ or caging; and iv/ widespread influence on gene expression. These features allow ParB to be classified as a Nucleoid-Associated Protein (NAP) involved in DNA compaction, genome topology maintenance and modulation of gene expression (77, 85, 86).

## SUPPLEMENTARY DATA

Supplementary Data are available at NAR Online.

## ACCESSION NUMBER

All sequencing data have been deposited at the Gene Expression Omnibus (GEO) under accession number GSE110158.

## ACKNOWLEDGEMENTS

ChIP DNA sequencing was performed by Genomed S.A. (Poland).

---

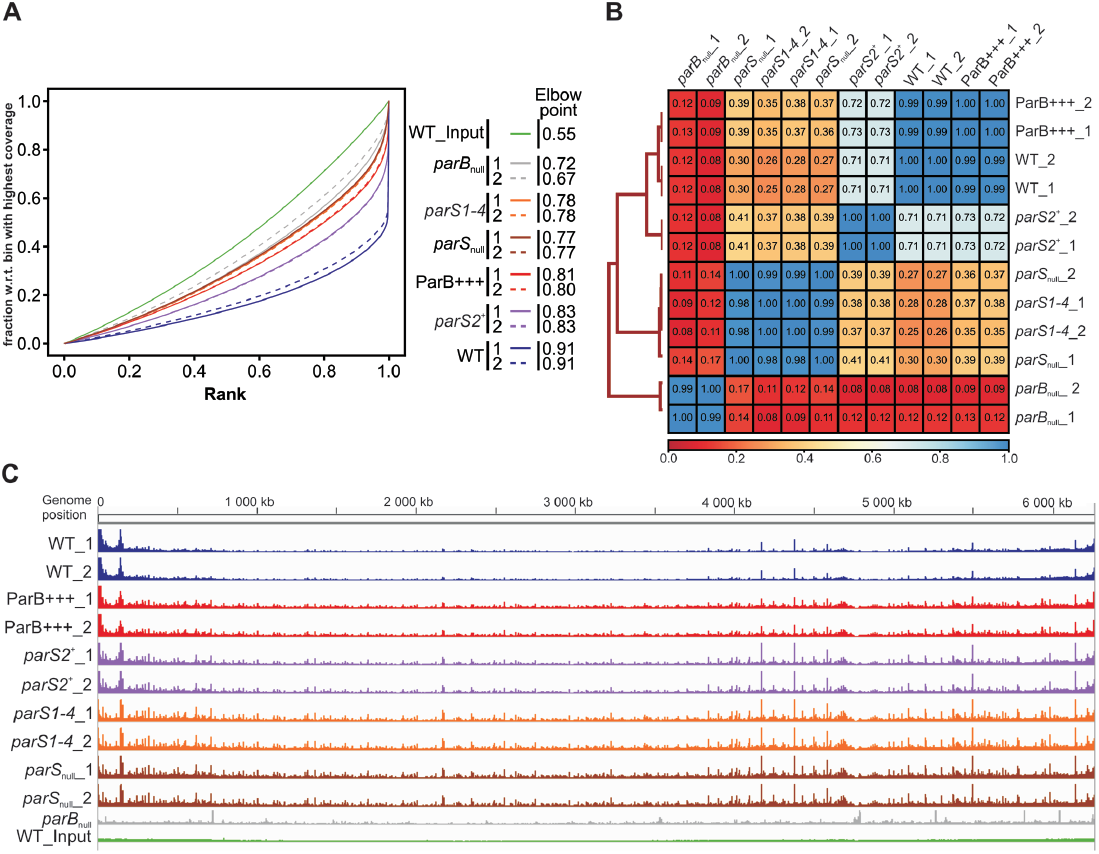

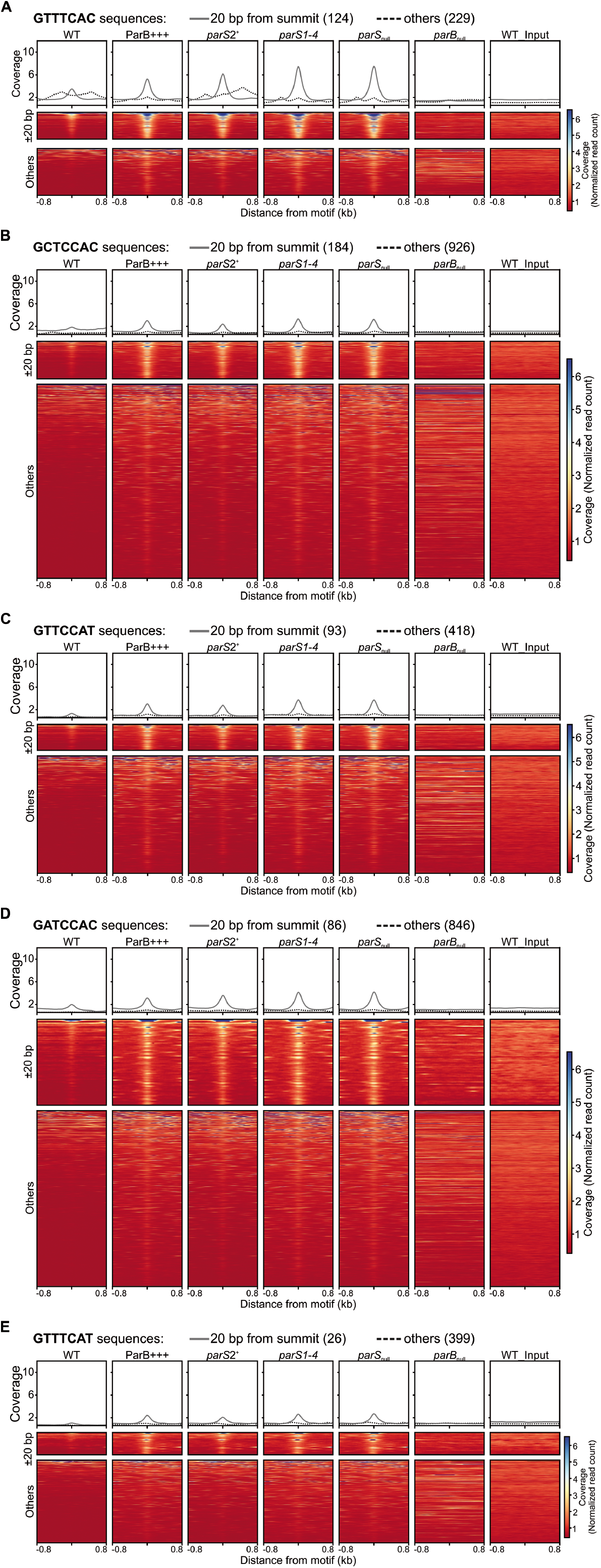

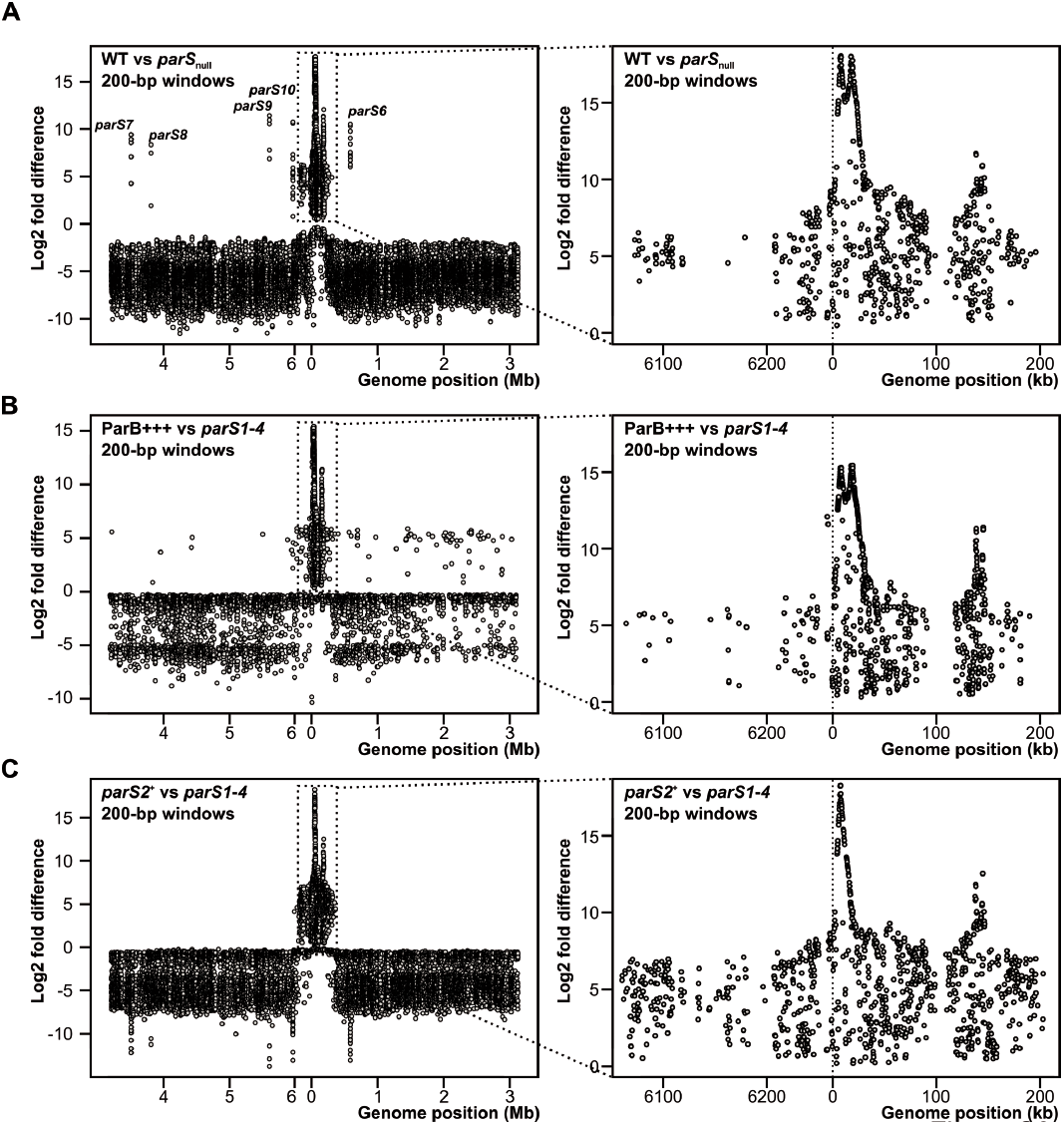

